# Gut microbiome signatures precede chronic kidney disease diagnosis in a large canine cohort

**DOI:** 10.64898/2026.08.25.746990

**Authors:** Hitomi Ono-Minagi, Nami Fujii, Masaki Ishikawa, Katsutoshi Tamura, Takayoshi Sakai

**Affiliations:** Advanced Research Center for Oral and Craniofacial Sciences, Okayama University Graduate School of Medicine, Dentistry and Pharmaceutical Sciences, Okayama, Japan; Department of Rehabilitation for Orofacial Disorders, Division of Growth and Development Dentistry, Graduate School of Dentistry, The University of Osaka, Suita, Japan; Anicom Big Data and MA-T Clinical Science Collaborative Course, Graduate School of Dentistry, The University of Osaka, Suita, Osaka, Japan; Department of Microbiology and Immunology, Institute for Immunological Sciences, Department of Orthopaedics, University of Rochester Medical Center, Rochester, NY, USA; Laboratory of Molecular Immunology, National Heart, Lung, and Blood Institute, National Institutes of Health, Bethesda, MD, USA; Anicom Specialty Medical Institute, Inc., Tokyo, Japan

**Keywords:** chronic kidney disease, gut microbiome, oral microbiome, companion dogs, longitudinal cohort

## Abstract

Chronic kidney disease (CKD)-associated dysbiosis is well described after diagnosis, but whether microbial changes precede clinical recognition is unclear. We integrated insurance claims, fecal and oral 16S rRNA profiles, and clinical laboratory data from companion dogs. Among 140,025 dogs, lower gut microbial diversity was associated with incident CKD after adjustment for age, sex and body size. Prediagnostic samples showed reduced evenness-related diversity, modest community shifts and seven differentially abundant genera. A five-genus score was elevated more than two years before diagnosis, although it was derived and evaluated in the same cohort and was not intended as a predictive model. In a laboratory subset, microbial changes preceded the largest increases in blood urea nitrogen and creatinine. Paired oral-gut samples showed limited exploratory associations between periodontal-associated taxa and the gut score. These findings identify microbial features associated with future claims-defined canine CKD and support independent validation and mechanistic investigation.

## INTRODUCTION

Chronic kidney disease (CKD) is a major public-health burden in humans and companion animals. Because early CKD is frequently asymptomatic, clinically recognized disease often reflects renal dysfunction that has already progressed beyond its earliest stages ^1^. Earlier biological signals associated with future CKD could improve understanding of disease development and may reveal modifiable pathways before overt clinical deterioration. Among candidate pathways, the microbiome is particularly attractive because it links mucosal ecology with host metabolism, inflammation and renal physiology ^2–4^.

In human CKD and experimental kidney disease, gut dysbiosis has been associated with altered microbial metabolism, impaired barrier function, systemic inflammation and the accumulation of uremic solutes ^2–4^. However, human CKD microbiome studies have largely focused on established disease, making it difficult to determine whether microbial disruption is already present during a preclinical period, reflects established renal dysfunction, or both. Longitudinal cohorts with microbiome samples collected before diagnosis are therefore needed to clarify the temporal relationship between mucosal microbial change and kidney disease ^5,6^.

Companion dogs provide a useful comparative setting for this question. They share household environments with humans, naturally develop age-associated diseases including CKD and periodontal disease, and receive longitudinal veterinary care that can be linked to large-scale health records ^7–11^. These features enable microbial profiles to be examined in relation to subsequent real-world clinical records, while naturally occurring canine CKD may also offer comparative insight relevant across species.

Oral health adds a second mucosal dimension to this framework. Periodontal disease is common in humans and companion dogs and has been linked to systemic inflammatory and cardiometabolic disease risk ^7,12,13^. Periodontal-associated bacteria can reach the gastrointestinal tract, where they may influence microbial ecology, mucosal immunity or barrier function ^14–16^. Together, these observations motivate investigation of a possible oral-gut-kidney connection, defined here as covariation among oral microbial burden, gut microbial ecology and renal health ^17^. However, paired oral-gut data linked to kidney outcomes remain limited, and observational associations cannot establish directionality or causality.

Here, we integrated insurance-claims records, fecal microbiome profiles, paired oral-gut samples and clinical laboratory measurements from companion dogs. Population-scale, matched case-control, longitudinal analyses showed that lower gut microbial diversity and a multi-genus signature were associated with claims-defined CKD before the first recorded diagnosis. A five-genus score summarized taxonomic differences across prediagnostic intervals and paired oral-gut samples revealed exploratory nominal associations between periodontal-associated oral taxa and the gut score. Together, these findings establish temporal associations that warrant independent validation and mechanistic studies of microbial changes accompanying future CKD claims.

## RESULTS

### Study cohorts for population-scale, matched, longitudinal and oral-gut analyses

We assembled four complementary cohorts to examine whether gut microbial features were associated with CKD before and after clinical diagnosis (Fig. 1 and Table 1). The population-scale discovery cohort comprised 140,025 dogs after sequence-quality control, including 1,215 dogs with a recorded CKD diagnosis. CKD ranked among the common claim reasons recorded in dogs whose policies ended because of death in companion animals in the source population, supporting its selection as a clinically relevant outcome (Supplementary Table 1). The matched case-control cohort included 756 dogs sampled before a subsequent CKD diagnosis (Future CKD), 400 dogs with CKD at sampling (Existing CKD), and age– and body-size-matched controls (Control F, n = 1,512; Control E, n = 800). Within the Future CKD group, 127 dogs had linked blood chemistry data available before diagnosis. The longitudinal cohort included 150 Future CKD dogs sampled twice and 300 matched controls; paired samples were separated by a mean of 318.4 ± 82.1 days (range, 265-762 days). The paired oral-gut cohort comprised 52 dogs with contemporaneous oral and fecal samples. Together, these cohorts supported population-level risk analysis, matched comparisons around diagnosis, within-dog longitudinal analysis, descriptive comparison of microbial and laboratory trajectories, and exploratory evaluation of oral-gut associations.

**Fig. 1.**
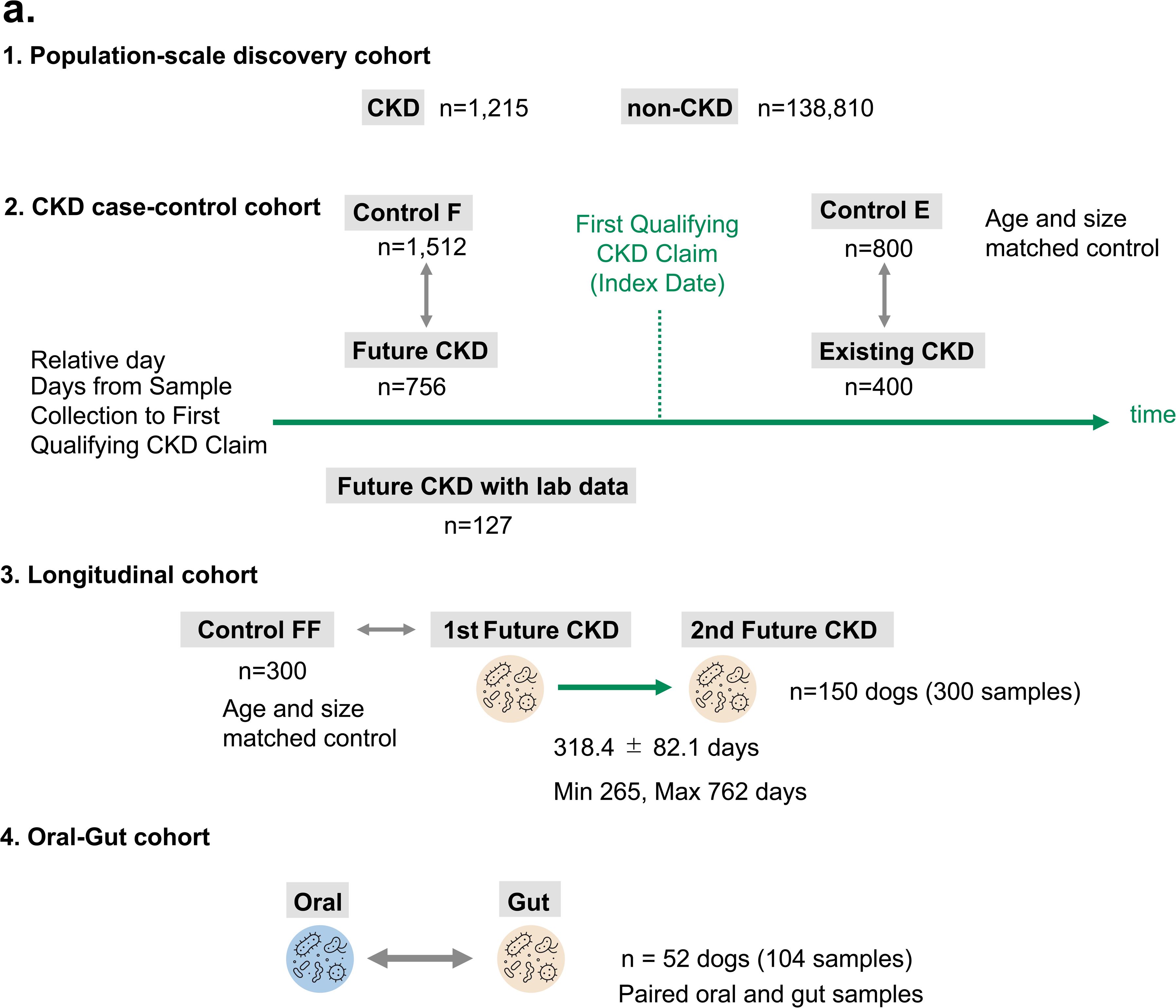
Study design and overview of the canine CKD microbiome cohorts. **a**. Schematic of the four analytical cohorts. The population-scale discovery cohort included 1,215 dogs with CKD and 138,810 dogs without CKD after quality control. The matched case-control cohort included dogs sampled before a subsequent CKD diagnosis (Future CKD, n = 756), dogs with CKD at sampling (Existing CKD, n = 400), and age– and body-size-matched controls (Control F, n = 1,512; Control E, n = 800). Relative day denotes the interval from microbiome sampling to the first qualifying CKD claim. A laboratory-data subset included 127 Future CKD dogs. The longitudinal cohort included 150 dogs sampled twice (300 samples) and 300 matched controls; the mean interval between paired samples was 318.4 ± 82.1 days (range, 265-762 days). The oral-gut cohort included 52 dogs with paired oral and fecal samples (104 samples). Table 1 summarizes cohort sizes and the cohort-specific sequence-processing parameters. CKD, chronic kidney disease; Control F, controls matched to Future CKD; Control E, controls matched to Existing CKD; Control FF, controls for the longitudinal cohort; FFCKD, longitudinal Future CKD samples; ASV, amplicon sequence variant; maxEE, maximum expected errors. **Supplementary** Table 1 Most frequently recorded reasons for insurance claims among deceased companion dogs and cats aged 0–7 years. Percentages were calculated separately for dogs (n = 424) and cats (n = 222) whose insurance policies were terminated because of death. The recorded conditions represent the reasons for insurance claims on the date of veterinary consultation and do not necessarily indicate the confirmed underlying causes of death.

**Table 1.**
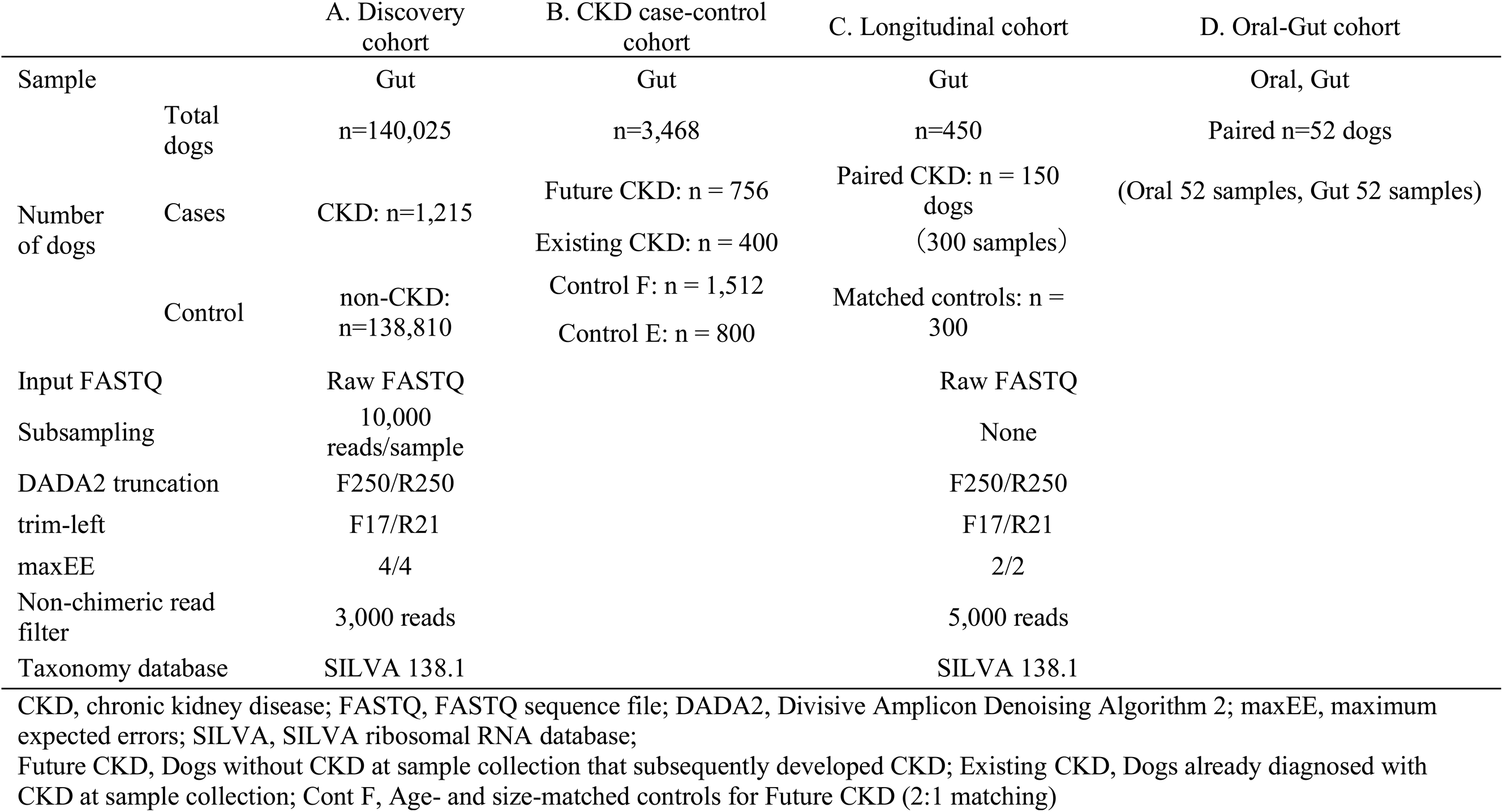
Overview of the study cohorts and microbiome analysis pipeline.

### Lower gut microbial diversity was associated with subsequent CKD diagnosis

Compared with dogs without CKD, dogs with CKD in the discovery cohort were older (6.71 ± 4.21 versus 2.97 ± 2.71 years) and differed in sex and body-size distributions (Supplementary Table 2). We first evaluated demographic covariates for subsequent microbiome analyses. In multivariable logistic regression, the odds of incident CKD increased with age (odds ratio [OR] per year, 1.374; 95% confidence interval [CI], 1.339-1.410; P = 2.23 × 10⁻¹³⁰) and were higher in medium-sized than in small dogs (OR, 1.692; 95% CI, 1.336-2.143; P = 1.29 × 10⁻⁵). Large body size and male sex were not independently associated with incident CKD in this model (Fig. 2a and Supplementary Table 3).

**Fig. 2.**
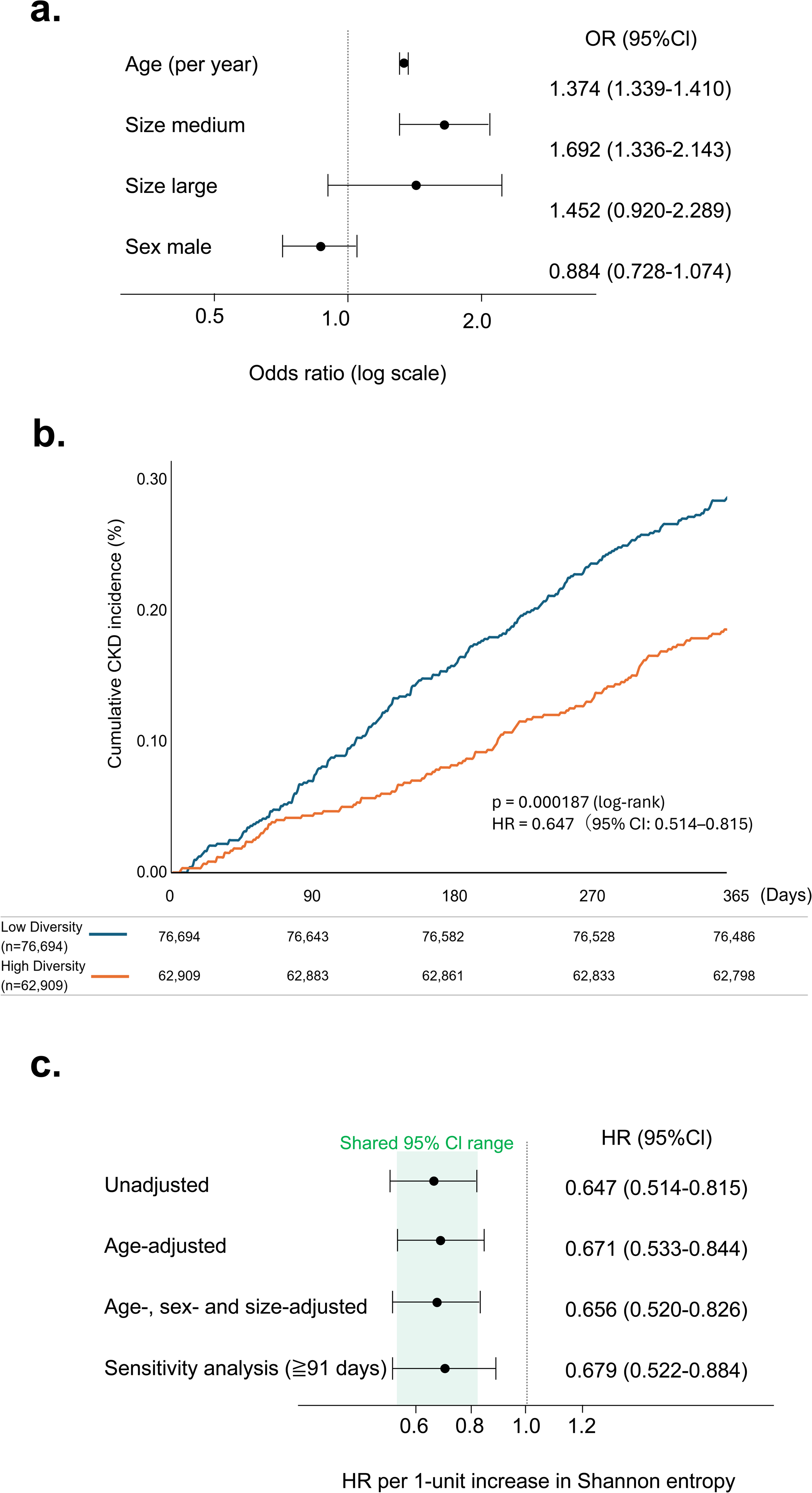
Lower gut microbial diversity precedes incident CKD in the population-scale cohort. **a**. Forest plot of the multivariable logistic-regression model for incident CKD, including age (per year), body size (medium or large versus small) and sex (male versus female). Points show ORs and horizontal lines show 95% CIs; the x axis is logarithmic. b. Cumulative 1-year incidence of CKD in dogs with low (n = 76,694) or high (n = 62,909) gut microbial diversity. Low and high diversity were defined using a pre-existing company classification based on breed– and age-standardized Shannon entropy scores, with cutoffs of 43 for small dogs, 45 for medium-sized dogs and 48 for large dogs. These cutoffs were established before the present study and were not selected using CKD outcomes. The inset gives the two-sided log-rank P value and the univariable Cox-model HR per one-unit increase in Shannon entropy (HR = 0.647, 95% CI 0.514–0.815; P = 0.00021). Numbers at risk are shown below the plot. c. Forest plot showing hazard ratios (HRs) for incident CKD per one-unit increase in Shannon entropy across the unadjusted, age-adjusted, age-, sex– and body-size-adjusted, and ≥91-day sensitivity Cox proportional-hazards models. Points indicate HRs and horizontal lines indicate 95% confidence intervals (CIs). The green shaded region indicates the range shared by the 95% CIs of all four models (HR, 0.533–0.815). Table 2 presents the results of univariable and multivariable Cox proportional hazards models, together with a sensitivity analysis restricted to dogs with ≥91 days of follow-up. Dogs without an available diversity measurement were excluded from the Cox analysis. **Supplementary** Table 2 summarizes the discovery cohort; sex and body-size distributions were compared using chi-square tests, and age and Shannon entropy using two-sided Welch t tests. **Supplementary** Table 3 gives multivariable logistic-regression results. OR, odds ratio; HR, hazard ratio; CI, confidence interval. *P < 0.05, **P < 0.01, ***P < 0.001; n.s., not significant.

We next asked whether gut microbial diversity was associated with CKD status and subsequent CKD diagnosis. Mean Shannon entropy was lower in dogs with CKD than in dogs without CKD (3.77 ± 0.71 versus 3.99 ± 0.72; P = 7.70 × 10⁻²⁶; Supplementary Table 2). Consistent with this cross-sectional difference, dogs in the low-diversity group had a higher cumulative 1-year incidence of CKD than dogs in the high-diversity group (log-rank P = 0.000187; Fig. 2b). When Shannon entropy was analyzed as a continuous predictor, each one-unit increase was associated with a 35.3% lower hazard of future CKD in the univariable Cox model (hazard ratio [HR], 0.647; 95% CI, 0.514-0.815; P = 0.00021). The association persisted after adjustment for age (HR, 0.671; 95% CI, 0.533-0.844; P = 0.00066) and after adjustment for age, sex and body size (HR, 0.656; 95% CI, 0.520-0.826; P = 0.00035). Excluding diagnoses made within 90 days of sampling yielded a similar estimate (HR, 0.679; 95% CI, 0.522-0.884; P = 0.0040), indicating that the association was not confined to dogs diagnosed shortly after sampling (Table 2 and Fig. 2c).

**Table 2.** Cox proportional hazards models for future CKD according to gut microbial diversity.

| Model |  | HR | 95% CI | p-value |  |
| --- | --- | --- | --- | --- | --- |
| Univariate |  | 0.647 | 0.514–0.815 | 0.00021 | *** |
| Multivariable analysis | Age-adjusted | 0.671 | 0.533–0.844 | 0.00066 | *** |
|  | Age + sex + size adjusted | 0.656 | 0.520–0.826 | 0.00035 | *** |
| Sensitivity analysis ($\geq 91$ days) | | 0.679 | 0.522–0.884 | 0.0040 | ** |

Hazard Ratio(HR), Confidence interval(CI)

### Gut microbial diversity and community structure differed across CKD groups

We next used the matched case-control cohort to examine whether the diversity signal observed in the population-scale analysis was accompanied by broader differences in gut microbiome structure. Shannon entropy differed among Control F, Future CKD and Existing CKD dogs (Kruskal-Wallis P = 0.002), as did Simpson diversity (P = 0.004; Fig. 3a). Both diversity indices were lower in Future CKD and Existing CKD dogs than in matched controls, whereas Future CKD and Existing CKD dogs did not differ from each other after correction for multiple comparisons. By contrast, neither observed features nor Chao1 richness differed significantly among groups (Kruskal-Wallis P = 0.084 and P = 0.083, respectively). These findings indicate that CKD-related differences were more apparent in diversity measures that incorporate taxon evenness than in richness alone. Comparable patterns were observed when Existing CKD dogs were compared with their separately matched controls (Control E).

**Fig. 3.**
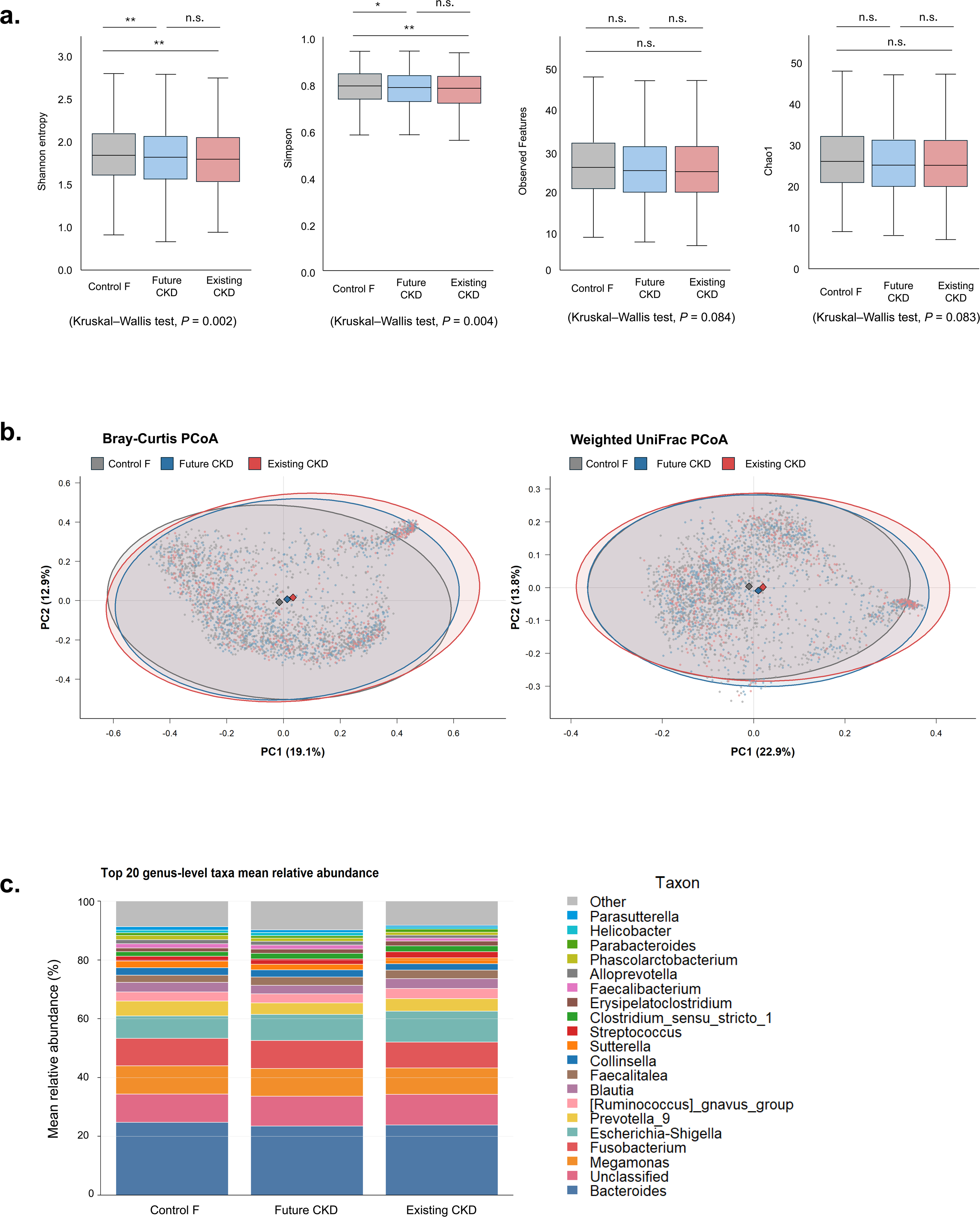
Gut microbial diversity and community structure differ across CKD groups. **a**. Shannon entropy, Simpson diversity, observed features and Chao1 richness in Control F (n = 1,512), Future CKD (n = 756) and Existing CKD (n = 400). Boxes show medians and interquartile ranges; whiskers extend to 1.5 times the interquartile range. Outliers are not displayed. Overall P values are from two-sided Kruskal-Wallis tests; pairwise comparisons used two-sided Wilcoxon rank-sum tests with BH correction. b. PCoA of Bray-Curtis (left) and weighted UniFrac (right) distances. Points represent individual dogs; shaded regions show 95% data ellipses and diamonds indicate group centroids. Overall PERMANOVA used 199 permutations (Bray-Curtis: pseudo-F = 3.728, R² = 0.00279, P = 0.005; weighted UniFrac: pseudo-F = 4.282, R² = 0.00320, P = 0.005). Pairwise BH-adjusted results are provided in **Supplementary** Table 4. c. Mean genus-level relative abundance of the 20 most abundant taxa in each group; remaining taxa are combined as Other. *q < 0.05, **q < 0.01; n.s., not significant.

Community composition also differed across the three groups. PERMANOVA detected overall differences in Bray-Curtis dissimilarity (pseudo-F = 3.728, R² = 0.00279, P = 0.005) and weighted UniFrac distance (pseudo-F = 4.282, R² = 0.00320, P = 0.005; Fig. 3b). In pairwise analyses, Control F differed from both Future CKD and Existing CKD for each distance metric (BH-adjusted q = 0.0075), whereas Future CKD and Existing CKD did not differ (Bray-Curtis q = 0.255; weighted UniFrac q = 0.23; Supplementary Table 4). Although statistically detectable, these shifts were small: group membership explained less than 0.3% of the variation in either distance metric. The mean abundance profiles of the 20 most abundant genera were broadly similar across groups, consistent with the modest beta-diversity effect sizes (Fig. 3c).

### Seven genera were associated with future CKD

Genus-level testing between Future CKD and Control F identified seven taxa that remained significant after Benjamini-Hochberg correction (Fig. 4a and Supplementary Table 5). Prevotella 9, Blautia, Peptoclostridium and Sutterella were less abundant in Future CKD dogs, whereas Clostridium sensu stricto 1, Pseudomonas and Incertae Sedis were more abundant. The largest positive fold change was observed for Pseudomonas (log2 fold change, 2.166; q = 0.0443), although this reflected a low-abundance genus. The remaining effects were smaller in magnitude, with log2 fold changes ranging from –0.526 for Peptoclostridium to 0.404 for Incertae Sedis (q = 0.0345-0.0455).

**Fig. 4.**
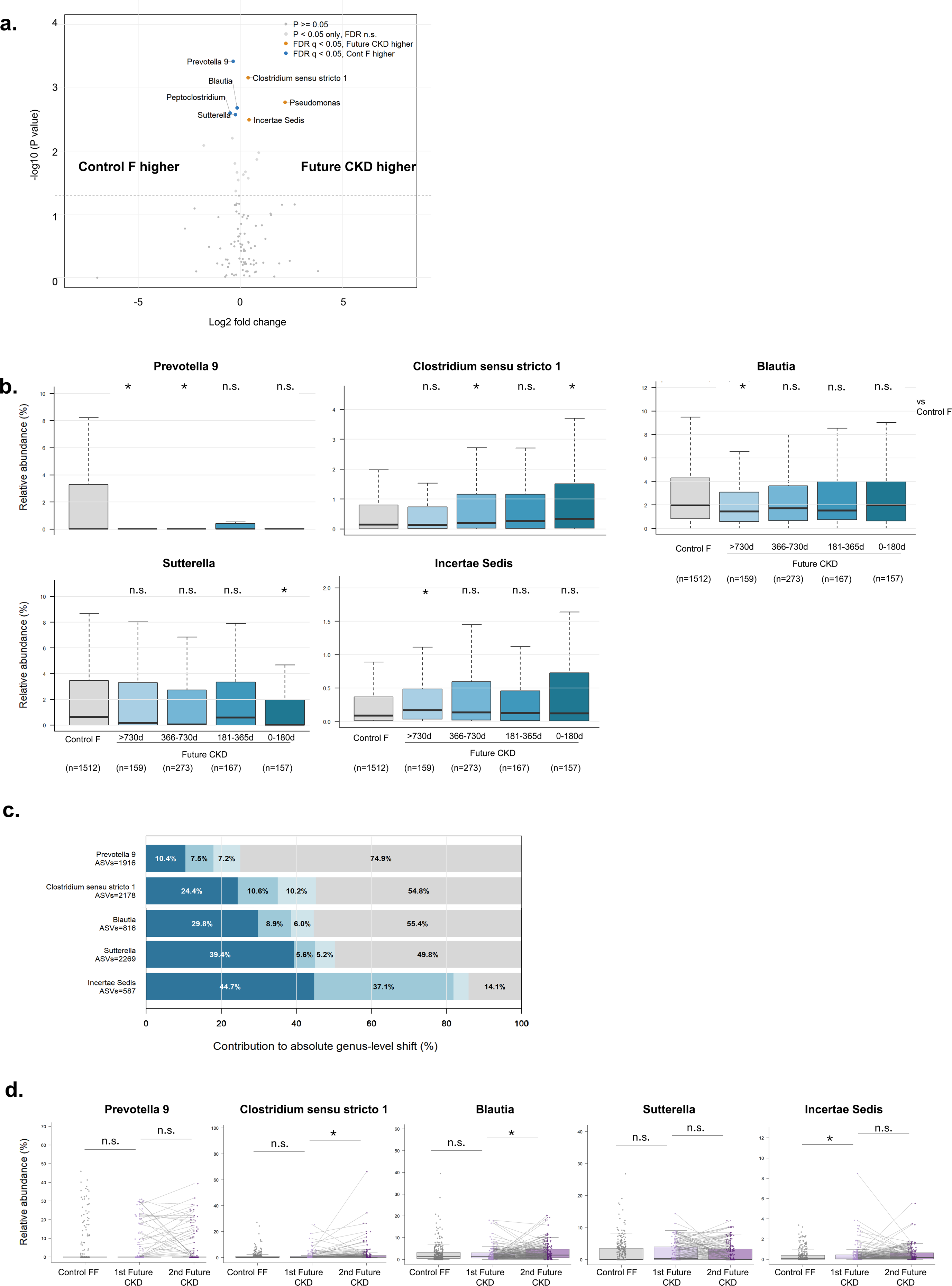
Genus-level signatures of future CKD and their temporal and longitudinal patterns. **a**. Differential genus abundance between Future CKD (n = 756) and Control F (n = 1,512). The x axis shows log2 fold change in mean relative abundance and the y axis −log10(two-sided Wilcoxon rank-sum P value). Orange and blue points indicate genera significantly enriched in Future CKD and Control F, respectively, after BH correction (q < 0.05); light-grey points indicate nominal P < 0.05 but q ≥ 0.05. Seven genera remained significant. Table 3 summarizes robustness across Wilcoxon, ANCOM-BC and covariate-adjusted MaAsLin2 analyses; Supplementary Table 5 lists the top 20 genera. b. Relative abundances of the five genera included in the gut CKD score in Control F and Future CKD dogs stratified by time to diagnosis (>730 days, n = 159; 366–730 days, n = 273; 181–365 days, n = 167; 0–180 days, n = 157). Boxes show medians and interquartile ranges; whiskers extend to 1.5× the interquartile range. Comparisons used two-sided Wilcoxon rank-sum tests with BH correction across 28 genus-by-time-bin comparisons. Pseudomonas and Peptoclostridium are shown in **Supplementary Fig. 1**. c. ASV-level decomposition of genus-level differences between Future CKD and Control F. The three ASVs contributing most strongly to each difference are shown separately; remaining ASVs are combined as “Other”. d. Longitudinal relative abundances in Control FF (n = 300) and paired first and second samples from 150 Future CKD dogs. Lines connect samples from the same dog. Control FF versus first Future CKD samples used two-sided Wilcoxon rank-sum tests; paired first versus second samples used two-sided Wilcoxon signed-rank tests. These comparisons were exploratory and are reported using nominal P values. Genus-wide longitudinal results are provided in **Supplementary** Table 6. In b, *q < 0.05 and **q < 0.01; in d, *P < 0.05 and **P < 0.01; n.s., not significant.

**Table 3.** Selection and robustness of seven genera associated with future CKD.

| Genus | log2FC <sup>1</sup> | Wilcoxon<br>q | Temporal<br>support <sup>2</sup> | ANCOM-BC |  | MaAsLin2 |  | Concordant<br>direction <sup>5</sup> |
| --- | --- | --- | --- | --- | --- | --- | --- | --- |
| | | | | logFC <sup>3</sup> | BC q | $\beta$ <sup>4</sup> | q | |
| Prevotella 9 | −0.379 | 0.0345 | Yes | −0.580 | 0.0507 | −0.915 | 0.0014 | ✓ |
| Clostridium sensu stricto 1 | 0.353 | 0.0345 | Yes | 0.365 | 0.0867 | 0.652 | 0.002 | ✓ |
| Blautia | −0.177 | 0.0443 | Yes | −0.291 | 0.0522 | −0.359 | 0.0112 | ✓ |
| Sutterella | −0.262 | 0.0443 | Yes | −0.522 | 0.0507 | −0.789 | 0.0051 | ✓ |
| Incertae Sedis | 0.404 | 0.0455 | Yes | 0.184 | 0.4126 | 0.326 | 0.0575 | ✓ |
| Pseudomonas | 2.166 | 0.0443 | Limited | 0.095 | 0.226 | 0.201 | 0.0023 | ✓ |
| Peptoclostridium | −0.526 | 0.0443 | Limited | −0.376 | 0.0507 | −0.450 | 0.0051 | ✓ |

<sup>1</sup> Descriptive log2 fold change calculated from the mean relative abundances in Future CKD versus Control F; statistical significance was evaluated using the Wilcoxon rank-sum test.
<sup>2</sup> Temporal support was assessed across four pre-diagnostic intervals (>730, 366–730, 181–365, and 0–180 days before CKD diagnosis).
<sup>3</sup> ANCOM-BC log fold change for Future CKD relative to Control F.
<sup>4</sup> MaAsLin2 coefficient for Future CKD relative to Control F, adjusted for age, sex, and body size. MaAsLin2 was applied to full-library TSS-normalized abundances using log transformation.
<sup>5</sup> Concordant direction indicates agreement in the direction of association across the descriptive log2 fold change, ANCOM-BC, and MaAsLin2 analyses.
P values were adjusted using the Benjamini–Hochberg method. Statistical significance was defined as $q < 0.05$ .

To assess the robustness of these associations to alternative analytical approaches, we re-analysed the same Future CKD (n = 756) and Control F (n = 1,512) samples using ANCOM-BC and covariate-adjusted MaAsLin2 (Table 3). All seven genera showed effect directions concordant with the original Wilcoxon analysis, but statistical support differed by method. None remained significant after genus-wide FDR correction in ANCOM-BC, although Prevotella 9, Peptoclostridium and Sutterella (q = 0.0507) and Blautia (q = 0.0522) were close to the prespecified q < 0.05 threshold. In MaAsLin2 models adjusted for age, sex and body size, six of the seven remained significant after standard multiple-testing correction; Incertae Sedis showed the same positive direction but did not reach the FDR threshold (q = 0.0575). Thus, directional consistency was observed across methods, whereas FDR-adjusted statistical significance was method-dependent.

Stratification by time to diagnosis showed that some components of this signature were already altered before the first CKD claim (Fig. 4b and Supplementary Fig. 1a). Compared with Control F, Prevotella 9 was reduced in the >730-day and 366–730-day groups, Blautia was reduced in the >730-day group, and Sutterella was reduced in the 0–180-day group. Clostridium sensu stricto 1 was increased in the 366–730-day and 0–180-day groups, whereas Incertae Sedis and Pseudomonas were increased in the >730-day group after correction across the genus-by-time comparisons. Other genus-by-time comparisons did not remain significant, indicating that individual taxa followed heterogeneous and non-monotonic trajectories. ASV-level decomposition further showed that the genus-level differences were not uniformly attributable to all sequence variants within a genus, but instead reflected contributions from selected ASVs, with remaining ASVs grouped as Other (Fig. 4c and Supplementary Fig. 1b).

We then examined the seven genera in a separate longitudinal cohort. At the first Future CKD sample, most genera did not differ from matched longitudinal controls (Control FF), although Incertae Sedis was higher at nominal P < 0.05 (Fig. 4d). Within the same dogs, Clostridium sensu stricto 1 and Blautia increased between the first and second samples (nominal P < 0.05 and P < 0.01, respectively). Prevotella 9, Sutterella, Incertae Sedis and Pseudomonas did not change significantly between paired samples. Because these seven-genus longitudinal comparisons were exploratory and were not adjusted for multiplicity, they describe within-dog temporal variation but do not confirm the behaviour of any individual taxon. In a complementary genus-wide longitudinal analysis of 99 eligible genera, Parasutterella decreased between the first and second samples (log2FC = −1.572, P = 3.18 × 10⁻⁵, q = 0.003), whereas Romboutsia increased (log2FC = 0.777, P = 0.00070, q = 0.035). No other genus remained significant after Benjamini–Hochberg correction (Supplementary Table 6).

### A five-genus CKD score was elevated more than two years before diagnosis

Because individual genera showed heterogeneous temporal patterns, we constructed a five-genus gut CKD score from the seven FDR-significant genera, retaining taxa with prediagnostic temporal support and sufficient abundance and directional consistency. The score included taxa increased in Future CKD dogs (Clostridium sensu stricto 1 and Incertae Sedis) and taxa reduced in Future CKD dogs (Prevotella 9, Blautia and Sutterella); higher values therefore indicate a more CKD-like profile within this dataset. In the same case-control cohort used both to select the component genera and evaluate the score, mean values were higher in Future CKD dogs than in Control F in every prediagnostic interval examined: >730 days (q < 0.01), 366-730 days (q < 0.001), 181-365 days (q < 0.05) and 0-180 days (q < 0.001; Fig. 5a). The pattern was not monotonic, although the largest mean difference occurred in the 0-180-day interval. The score was not intended as a predictive model, and these within-cohort analyses do not estimate out-of-sample discrimination, calibration or clinical utility.

**Fig. 5.**
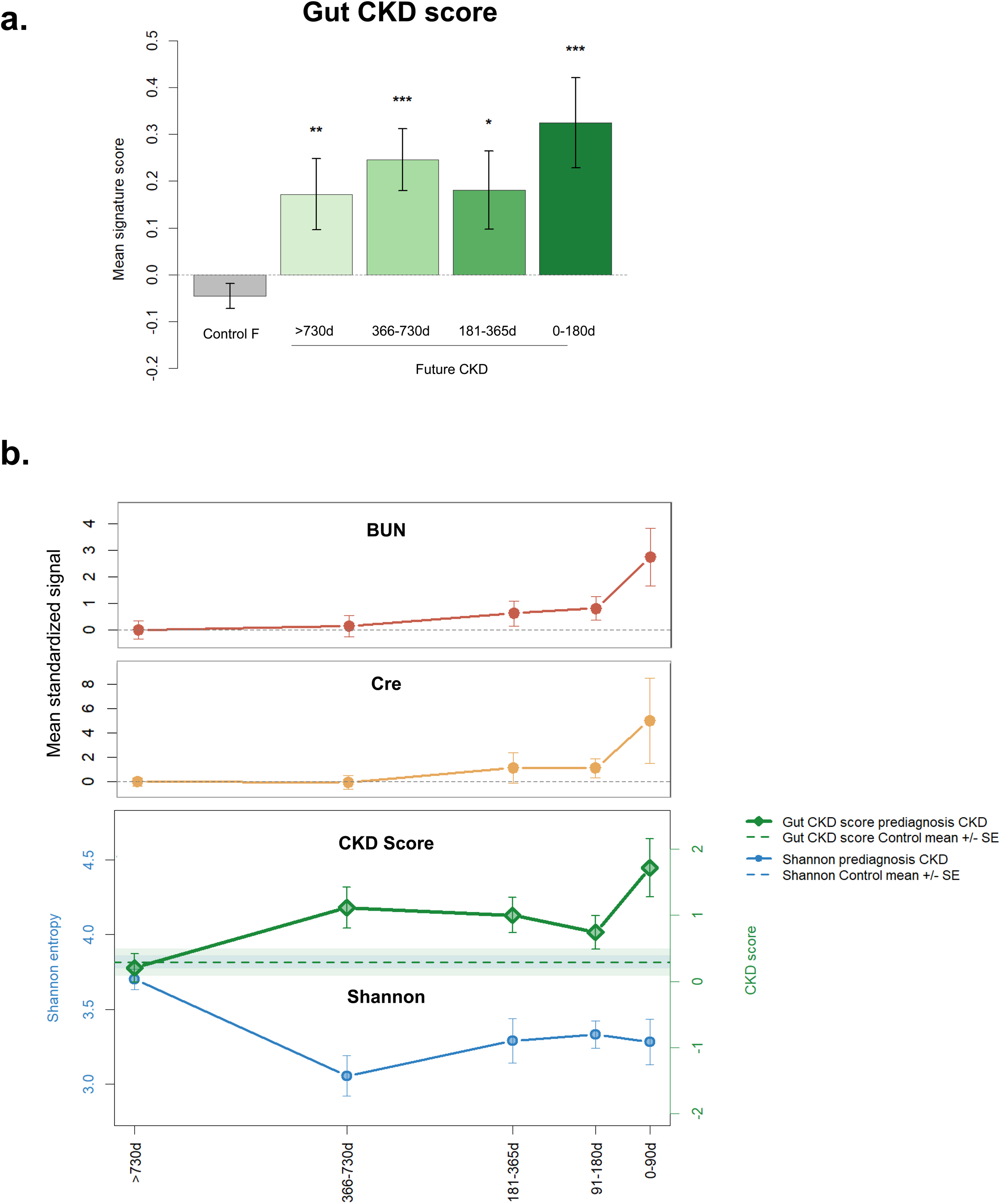
A gut microbial CKD score is elevated before clinical diagnosis. **a**. Mean five-genus gut CKD score in Control F (n = 1,512) and Future CKD dogs stratified by time to diagnosis (>730 days, n = 159; 366-730 days, n = 273; 181-365 days, n = 167; 0-180 days, n = 157). Bars and error bars show mean ± s.e.m. The score is the mean control-standardized log-abundance z score of Clostridium sensu stricto 1 and Incertae Sedis minus the mean corresponding z score of Prevotella 9, Blautia and Sutterella. Each time bin was compared with Control F using a two-sided Wilcoxon rank-sum test with BH correction. The score was derived and evaluated in this case-control cohort. b. Prediagnostic trajectories in the laboratory-data subset (127 Future CKD dogs). BUN and creatinine (top) were standardized using measurements obtained >730 days before diagnosis. The lower panel shows mean Shannon entropy (blue, left axis) and mean gut CKD score (green, right axis); dashed lines and shaded bands show matched-control means ± s.e.m. Points and error bars show mean ± s.e.m.; the number of available dogs varies by analyte and time bin and is provided in Source Data. *q < 0.05, **q < 0.01, ***q < 0.001 versus Control F; BUN, blood urea nitrogen; Cre, creatinine.

We descriptively compared the microbial pattern with prediagnostic laboratory trajectories in the 127 Future CKD dogs with linked measurements (Fig. 5b). BUN and creatinine remained near the distant prediagnostic reference in earlier intervals and increased most prominently during the 0-90 days preceding diagnosis. By contrast, lower Shannon entropy and a higher gut CKD score were apparent in earlier prediagnostic intervals, with the largest Shannon difference at 366-730 days before diagnosis. The number of dogs differed by analyte and time bin, and microbiome and laboratory measurements were aligned separately to the first CKD claim. These trajectories therefore describe timing and do not show that the microbial score independently predicts laboratory deterioration.

### Periodontal-associated oral taxa showed exploratory associations with the gut CKD score

We next examined paired oral and gut samples from 52 dogs to explore whether periodontal-associated oral taxa were linked to the gut CKD signature. Oral samples were collected from periodontal sites as illustrated on Fig. 6a. Oral and gut communities separated clearly by ASV-level Bray-Curtis dissimilarity (Fig. 6b). Paired PERMANOVA yielded pseudo-F = 17.06, R² = 0.143 and P = 0.001, with permutations restricted within dogs. This site effect provided the context for testing covariation between periodontal-associated oral taxa and the gut CKD signature rather than direct overlap between oral and gut communities.

**Fig. 6.**
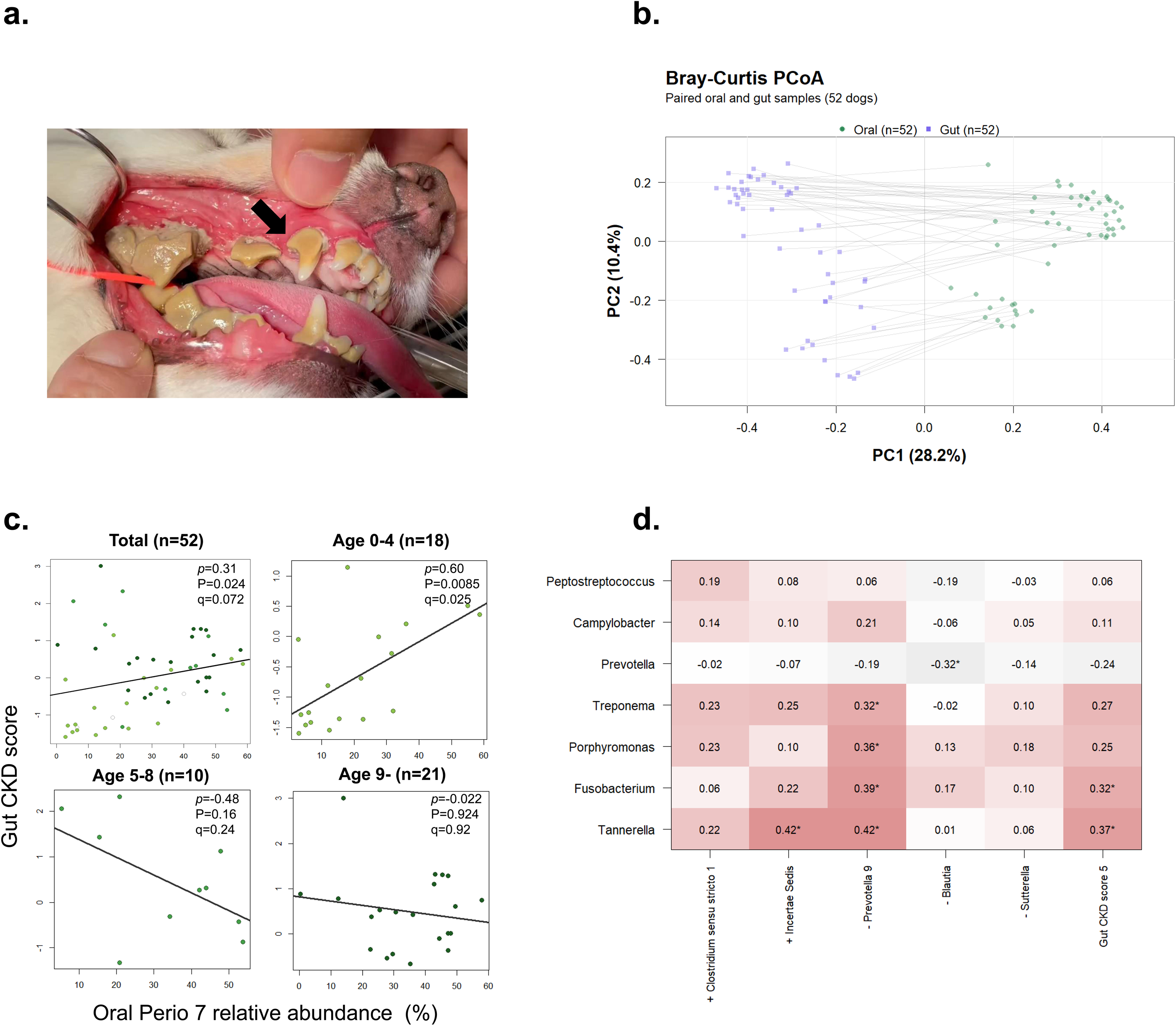
Exploratory associations between oral periodontal-associated taxa and the gut CKD score. **a**. Representative image of the canine oral sampling site. Arrows indicate the gingival margin and subgingival plaque. b. ASV-level Bray-Curtis PCoA of paired oral (green circles) and gut (purple squares) samples from 52 dogs. Grey lines connect samples from the same dog. Axis labels give the variance explained. Site separation was tested by paired PERMANOVA with 999 permutations restricted within dogs (pseudo-F = 17.06, R² = 0.143, P = 0.001). c. Exploratory association between Oral Perio7, the summed oral relative abundance of Porphyromonas, Treponema, Tannerella, Fusobacterium, Prevotella, Campylobacter and Peptostreptococcus, and the five-genus gut CKD score in all dogs and within age strata. Points represent dogs, and lines show least-squares fits for visualization; ρ and P are two-sided Spearman statistics. Total: n = 52, ρ = 0.313, P = 0.024; age 0-4 years: n = 18, ρ = 0.600, P = 0.0085; age 5-8 years: n = 10, ρ = –0.479, P = 0.162; age ≥9 years: n = 21, ρ = –0.022, P = 0.924. These unadjusted age-stratified analyses are exploratory and do not establish an age interaction; three dogs with missing age were excluded only from the stratified analyses. d. Spearman correlation matrix between the seven oral periodontal-associated genera (rows) and the five gut-score components and composite score (columns). Positive-direction gut genera are marked with + and negative-direction genera with –. Cell values are Spearman ρ. *P < 0.05 (two-sided, unadjusted exploratory results). **Supplementary** Table 7 summarizes the demographic characteristics of the oral–gut cohort, including sample size, sex distribution, body size, and age.

For this exploratory analysis, we defined Oral Perio7 as the summed oral relative abundance of seven periodontal-associated genera: Porphyromonas, Treponema, Tannerella, Fusobacterium, Prevotella, Campylobacter and Peptostreptococcus ^12,13,15^. Across all 52 dogs, Oral Perio7 showed a nominal positive correlation with the gut CKD score (Spearman ρ = 0.313, P = 0.024; Fig. 6c). The nominal association was stronger in dogs aged 0-4 years (n = 18, ρ = 0.600, P = 0.0085) but was not detected in dogs aged 5-8 years (n = 10, ρ = –0.479, P = 0.162) or ≥9 years (n = 21, ρ = – 0.022, P = 0.924). At the genus level, nominal correlations link Tannerella, Fusobacterium, Porphyromonas and Treponema with individual score components, and Tannerella and Fusobacterium with the composite score (Fig. 6d). These unadjusted correlations were based on a small paired cohort and small age strata and do not establish an age interaction, an oral-gut-kidney axis or a causal oral-gut pathway. Moreover, the overall association was attenuated after adjustment for age and sex (partial Spearman ρ = 0.055, 95% CI, −0.236 to 0.336; P = 0.710). Accordingly, these observations should be regarded as preliminary and hypothesis-generating.

## DISCUSSION

This study used naturally occurring CKD in companion dogs to examine whether microbial changes are detectable before clinical recognition of kidney disease. Across population-scale, matched and longitudinal analyses, lower gut microbial diversity and a multi-genus signature were associated with CKD before the first recorded claim, including in samples collected more than two years before diagnosis. Paired oral-gut samples also revealed nominal exploratory associations between periodontal-associated oral taxa and the gut CKD score. These data therefore establish temporal associations with future claims-defined CKD, but do not demonstrate that the microbial features cause CKD or predict it in new populations.

Confidence in this temporal association is strengthened by convergence across population-scale risk analysis, alignment to future diagnosis, matched case-control comparison and within-dog longitudinal sampling. In the discovery cohort, lower Shannon entropy was associated with incident CKD after adjustment for age, sex and body size. In the matched cohort, Future CKD dogs showed reduced evenness-related diversity and modest community shifts. The beta-diversity effect sizes were small, with CKD group explaining less than 0.3% of microbial variation in this heterogeneous real-world population. Thus, the principal signal is a modest population-level association rather than a discrete CKD-specific community state. Nevertheless, a small effect on overall community composition does not exclude biological relevance of particular taxa or microbial functions, because targeted changes in metabolite production or host-microbe interactions may occur without a large global shift in beta diversity.

The five-genus CKD score compactly summarized the selected taxonomic pattern, but the component genera were selected, and the score was evaluated in the same case-control cohort. Consequently, the score is descriptive and was not intended as a predictive model. Its elevation across prediagnostic intervals shows that the selected pattern was detectable before the first CKD claim but does not estimate out-of-sample discrimination, calibration or clinical utility. Independent prospective cohorts are required to evaluate reproducibility and any potential value for risk stratification.

Although periodontal disease has previously been associated with an increased risk of subsequent CKD in dogs^17^, the microbial links connecting the oral cavity, gut and kidney remain poorly understood. The paired oral-gut analysis provides preliminary, hypothesis-generating evidence of covariation between periodontal-associated oral taxa and the gut CKD score. Oral and gut communities were distinct, and the observed associations were limited and attenuated after adjustment for age and sex. Several non-exclusive explanations remain possible: repeated swallowing may introduce oral taxa into the gut, periodontal inflammation may influence systemic immune tone, or dysbiosis at both sites may reflect shared host or environmental factors. However, the small cross-sectional cohort cannot resolve directionality, age-specific effects or causality and therefore does not establish an oral-gut-kidney axis. Larger paired oral-gut cohorts, longitudinal sampling and mechanistic studies will be required to determine whether oral microbial alterations contribute to gut dysbiosis or CKD progression.

These findings are consistent with emerging human concepts linking mucosal immunity, periodontal inflammation and kidney disease. Gut dysbiosis has been proposed to promote systemic inflammation, barrier dysfunction and altered microbial metabolite production in CKD ^2–4^. Even modest alterations in particular taxa could be biologically relevant if they alter the production of microbial metabolites, including precursors of uremic solutes, or modulate mucosal and systemic immune responses ^2–4^. Periodontal disease has also been associated with systemic inflammatory burden and CKD risk ^12,18^. In addition, mucosal immune pathways, including IgA production, provide a plausible biological bridge between chronic microbial stimulation and renal vulnerability ^19–21^. The present study did not measure microbial metabolites, inflammatory mediators, IgA responses or renal histopathology; therefore, the functional implications of the identified taxa and these proposed mechanisms remain to be established. The temporal microbial associations observed here provide a rationale for future studies integrating metagenomics, metabolomics and immunological measurements to test these pathways directly.

Several limitations should guide interpretation. Claims-defined CKD reflects real-world veterinary practice but was not centrally adjudicated using uniform laboratory criteria or International Renal Interest Society staging. Diagnostic practices and the timing of claims may have varied among clinics, and records classified as suspected disease were included. Consequently, some cases of acute kidney injury or other renal disorders may have been misclassified as CKD. The first qualifying claim therefore represents the first recorded recognition of renal disease rather than its biological onset, and variation in the interval between these events may have obscured temporal relationships. In addition, some dogs contributed to more than one of the population-scale, case-control and longitudinal cohorts. Although each analysis addressed a distinct question, convergence across these cohorts should not be regarded as fully independent replication; the oral– gut cohort, however, was independent. Residual confounding by breed, diet, medication use, comorbidities, geographic region, sampling period and other environmental factors also remains possible. Furthermore, 16S rRNA gene sequencing limits strain-level resolution and functional interpretation, while the five-genus score was derived and evaluated in the same cohort and requires external validation before any predictive or clinical interpretation. Laboratory data were available for only a subset of dogs, and the oral–gut cohort was small and cross-sectional. Collectively, these limitations constrain causal and predictive inference and highlight the need for prospective studies incorporating standardized renal phenotyping, independent cohorts, longitudinal sampling and functional measurements.

In summary, this study identifies temporal associations between prediagnostic gut microbial features and future claims-defined CKD in companion dogs and reports preliminary associations between periodontal-associated oral taxa and the gut CKD signature. These observations provide a comparative framework for investigating a possible oral-gut-kidney connection. Independent prospective validation and mechanistic studies are required to determine whether the observed signatures reflect biologically relevant pathways and whether they have any value for risk assessment or prevention.

## METHODS

### Study design and data sources

We conducted a retrospective observational study with longitudinal components, integrating insurance-claims records, fecal and oral 16S rRNA gene amplicon profiles and, for a subset of dogs, clinical laboratory measurements. The primary dataset was a population-scale discovery cohort. Three additional analytical cohorts were assembled: (1) an age– and body-size-matched CKD case-control cohort; (2) a longitudinal cohort of dogs with repeated fecal samples; and (3) a paired oral-gut cohort. The unit of analysis was the individual dog except in explicitly paired longitudinal and oral-gut analyses. Investigators were not blinded to cohort labels during retrospective computational analyses. No prospective sample-size calculation was performed; all samples meeting predefined eligibility and sequence-quality criteria were included.

### Ethics and regulatory oversight

All procedures complied with relevant ethical regulations for animal research. The study protocol was reviewed and approved by the Animal Experiment Committee of Graduate School of Dentistry, The University of Osaka (approval no. R-07-035-0). The study included privately owned companion dogs enrolled in the Anicom insurance and health-management programme. A veterinarian explained the study to owners, who provided consent for the use of samples and associated clinical information. Fecal, oral and blood samples were collected during routine health examinations, and all data were de-identified before analysis. No animals were euthanized for this study.

### CKD ascertainment and clinical variables

CKD status was ascertained from insurance claims submitted by primary-care veterinary practices. Diagnoses were assigned by attending veterinarians during routine clinical care and recorded through the insurance claim process. Dogs were classified as having claims-defined CKD if at least one claim contained a prespecified label corresponding to chronic kidney disease, renal disease or renal failure, including labels recorded as suspected disease. The date of the first qualifying CKD-related claim was defined as the index date. Diagnoses were not independently adjudicated using standardized laboratory criteria or International Renal Interest Society staging; therefore, the index date represents the first recorded claim rather than the biological onset of CKD, and misclassification with acute kidney injury or other renal disorders cannot be excluded. Dogs with a qualifying claim on or before microbiome sampling constituted the Existing CKD group, whereas dogs without a qualifying claim at sampling that subsequently received one constituted the Future CKD group. These capitalized terms denote the prespecified analytical groups; “prediagnostic” refers descriptively to samples obtained before the first qualifying claim. The interval to diagnosis was calculated from microbiome sample collection to that claim. Dogs without a qualifying renal disease claim during available follow-up were classified as non-CKD controls, subject to cohort-specific eligibility criteria. Age at sampling, sex and body-size category were obtained from the insurance database. Body size was categorized by recorded body weight as small (<10 kg), medium (10 to <20 kg) or large (≥20 kg). For Future CKD dogs with clinical laboratory data, BUN and serum creatinine measurements were aligned to the interval between examination and the first qualifying claim; microbiome measurements were aligned independently to the interval between fecal sampling and that claim.

### Cohort overview

#### Population-scale discovery cohort

The discovery cohort comprised 140,025 dogs after quality control, including 1,215 dogs with CKD and 138,810 dogs without CKD. This cohort was used to examine population-level associations of demographic variables and gut microbial alpha diversity with CKD and incident CKD.

#### CKD case-control cohort

The CKD case-control cohort comprised of 756 Future CKD dogs, 400 Existing CKD dogs, 1,512 controls matched to Future CKD dogs (Control F) and 800 controls matched to Existing CKD dogs (Control E). Controls were selected at a 2:1 ratio from dogs without CKD and matched separately to the relevant case group using the age and body-size categories specified in the prespecified matched-list file. Because the primary objective of the present study was to characterize prediagnostic microbiome changes, subsequent genus-level differential-abundance and CKD score analyses focused on Future CKD and Control F; Control E was retained for cohort definition but was not included in these primary prediagnostic analyses.

#### Longitudinal cohort

The longitudinal cohort included 150 dogs that contributed two fecal samples each (1st and 2nd Future CKD; 300 samples total) and 300 age– and size-matched controls (Control FF). The mean interval between paired samples was 318.4 ± 82.1 days (range, 265-762 days).

#### Paired oral-gut cohort

The oral-gut cohort included 52 dogs with paired oral and fecal samples. The cohort included 25 male and 27 female dogs and had a mean age of 6.76 ± 5.54 years. Body-size information was available for 46 dogs (32 small, 10 medium and 4 large); size was unknown for 6 dogs. Age-stratified analyses used 0-4, 5-8 and ≥9 years; dogs with missing age were excluded only from the stratified analyses.

### Sample collection and storage

Fecal samples were collected using a dedicated collection kit containing a DNA stabilization solution (product no. 25-3606-H; Sugiyama-Gen Co., Ltd., Japan), transported to the laboratory at ambient temperature by standard mail, and generally received within several days to approximately 1 week. Upon arrival, samples were stored frozen until DNA extraction and thawed immediately before processing. Oral samples were collected by swabbing dental plaque from the canine tooth region using either a conventional cotton swab or the proprietary TF04 collection kit. The swab was immersed in the same DNA stabilization solution used for fecal samples, and oral samples were transported and stored under the same conditions as the fecal samples.

### DNA extraction, amplification and sequencing

Microbial DNA was extracted from fecal and oral samples at Anicom Specialty Medical Institute Inc. using the CMG-1076 chemagic DNA Stool 200 Kit H96 (Revvity chemagen Technologie GmbH, Baesweiler, Germany) on a chemagic 360 automated extraction system (Revvity chemagen Technologie GmbH), according to the manufacturer’s instructions. Extracted DNA was quantified using the Qubit dsDNA Quantification Assay (Thermo Fisher Scientific, Waltham, MA, USA).

The V3–V4 region of the bacterial 16S rRNA gene was amplified using the forward primer 5′-(TCGTCGGCAGCGTCAGATGTGTATAAGAGACAG)CCTACGGGNGGCWGCAG-3′ and reverse primer 5′– (GTCTCGTGGGCTCGGAGATGTGTATAAGAGACAG)GACTACHVGGGTATCTAATCC-3′. Amplicon PCR was performed using a KAPA amplification kit (Kapa Biosystems, Wilmington, MA, USA), with an initial denaturation at 95 °C for 3 min, followed by 30 cycles of 95 °C for 30 s, 55 °C for 30 s and 72 °C for 30 s, and a final extension at 72 °C for 5 min. PCR products were purified using Sera-Mag beads and quantified before index PCR. Index PCR was performed using Nextera-compatible primers (Integrated DNA Technologies, Coralville, IA, USA) for 12 cycles under the same cycling temperatures.

Libraries were prepared using a protocol based on the Illumina 16S Metagenomic Sequencing Library Preparation workflow and pooled at equimolar concentrations. Sequencing was performed on Illumina platforms (Illumina, San Diego, CA, USA). Samples from Test No. 25 were sequenced on a MiSeq using the MiSeq Reagent Kit v3 with 2 × 300-bp paired-end reads, whereas samples from Test No. 64 were sequenced on a NovaSeq 6000 using the NovaSeq 6000 SP Reagent Kit v1.5 (500 cycles) and NovaSeq XP 2-Lane Kit v1.5 with 2 × 250-bp paired-end reads.

Laboratory quality control was conducted at the DNA extraction, first-PCR and second-PCR stages using standard (STD) and negative-control (NC) samples with predefined concentration criteria.

If an NC failed the relevant criterion, samples were generally reprocessed beginning with the DNA extraction step. Following sequencing, STD samples were evaluated for both taxonomic composition and Shannon entropy. If the cumulative relative abundance of the major expected taxa in an STD was <80%, the corresponding plate was considered potentially contaminated and classified as a QC failure. STD Shannon entropy was required to be within the predefined range of 4.4–5.5. Sample-level quality control was additionally performed by assessing the relative abundance of environmental bacterial taxa, and samples in which these taxa accounted for >70% of the detected community were excluded from downstream analyses.

### Amplicon sequence processing and taxonomic assignment

Sequence data were processed using QIIME 2 (version 2024.10). Paired-end reads were denoised with DADA2 using independent pooling and consensus-based chimera removal. Processing parameters differed among cohorts and are summarized in Table 1. For the case-control and longitudinal and oral–gut workflows, forward and reverse reads were truncated at 250 bases after removal of 17 and 21 bases from the 5′ ends of the forward and reverse reads, respectively, and a maximum expected-error threshold of 2 was applied to each read.

For the population-scale discovery cohort, each FASTQ file was randomly subsampled to 10,000 reads using seqtk before QIIME 2 processing. Maximum expected-error thresholds of 4 were applied to both forward and reverse reads, and samples yielding fewer than 3,000 non-chimeric reads were excluded. The case-control, longitudinal and oral–gut workflows were processed without prior subsampling and required at least 5,000 non-chimeric reads per sample.

Amplicon sequence variants (ASVs) were taxonomically assigned using the QIIME 2 classify-sklearn method with the SILVA reference database, release 138.1. Sequences assigned to mitochondrial, chloroplast or non-bacterial taxa were removed before downstream analysis.

For the case-control analyses, a precomputed genus-level abundance table was used. Genus-level taxonomic labels were generated by extracting the genus field from the taxonomic assignment. Missing or empty genus annotations were designated “Unclassified”, underscores in taxonomic labels were replaced with spaces, and features assigned to the same genus were aggregated. “Unclassified” was retained as an explicit category.

Within-sample diversity was evaluated using Shannon entropy, Simpson diversity, observed features and Chao1 richness. The primary case-control analysis included Control F (n = 1,512), Future CKD (n = 756) and Existing CKD (n = 400). Overall differences among groups were assessed using Kruskal–Wallis tests, followed by two-sided Wilcoxon rank-sum tests for pairwise comparisons, with P values adjusted for multiple comparisons using the Benjamini–Hochberg method. Genus-level community composition was summarized using the mean relative abundance of the 20 most abundant genera in each group, with all remaining taxa combined into an “Other” category.

### Beta diversity

For weighted UniFrac analysis, ASVs with a total frequency <500 across the analysed dataset were removed before phylogenetic reconstruction. Representative ASV sequences were aligned with MAFFT; alignment columns were masked using a minimum conservation threshold of 0.4 and a maximum gap frequency of 1.0; and an approximately maximum-likelihood tree was inferred with FastTree using QIIME 2’s align-to-tree-mafft-fasttree pipeline. The resulting rooted tree was used to calculate weighted UniFrac distances. Principal coordinates analysis (PCoA) was performed on the exported distance matrix for visualization. Ellipses in the three-group plots show 95% data ellipses, and diamonds indicate group centroids. Group differences were tested by one-way permutational multivariate analysis of variance (PERMANOVA) using 199 permutations in the three-group case-control analysis. Pairwise PERMANOVA P values were BH-adjusted. In the oral-gut cohort, site differences were tested using 999 permutations restricted within dogs to preserve pairing.

### Differential-abundance and diagnosis-timing analyses

For each genus, relative abundance in Future CKD and Control F was compared using a two-sided Wilcoxon rank-sum test. Genera were retained if they were detected (relative abundance >0) in at least 5% of samples in the combined Future CKD and Control F dataset or if their mean relative abundance in the combined dataset was at least 0.0001 (0.01%). Log2 fold change was calculated from group mean relative abundances using a pseudocount of 1 × 10⁻⁶, and P values were adjusted across tested genera using the Benjamini-Hochberg (BH) method. Seven genera had BH-adjusted q < 0.05: Prevotella 9, Clostridium sensu stricto 1, Pseudomonas, Blautia, Peptoclostridium, Sutterella and Incertae Sedis. For temporal analyses, Future CKD dogs were grouped by the interval from sampling to diagnosis (>730, 366-730, 181-365 or 0-180 days), and each bin was compared with Control F using two-sided Wilcoxon rank-sum tests. BH adjustment was applied across the 28 genus-by-time-bin comparisons. Overall differences across Control F and the four-time bins were assessed using Kruskal-Wallis tests.

### Sensitivity differential-abundance analyses

To assess the robustness of the seven genera identified in the primary Wilcoxon analysis, the same Future CKD (n = 756) and Control F (n = 1,512) samples were re-analysed using ANCOM-BC and MaAsLin2. ANCOM-BC was used to estimate bias-corrected log fold changes between Future CKD and Control F. MaAsLin2 was applied to full-library total-sum-scaled relative abundances after log transformation, with CKD group as the variable of interest and age, sex and body-size category included as covariates. P values were adjusted for multiple testing across the tested genera using the Benjamini–Hochberg procedure, with q < 0.05 considered statistically significant. Directional concordance was evaluated by comparing the signs of the effect estimates from the Wilcoxon, ANCOM-BC and MaAsLin2 analyses.

### ASV-level decomposition of genus shifts

For each CKD-associated genus, ASV relative abundances were averaged within Future CKD and Control F. The contribution of each ASV to the genus-level shift was calculated as the absolute difference between its mean relative abundance in Future CKD and Control F divided by the sum of absolute ASV-level differences within that genus. The three ASVs with the largest contributions were displayed separately, and all remaining ASVs were combined as “Other”. This decomposition describes the internal composition of the observed genus-level difference and was not used as an independent significance test.

### Longitudinal validation

Relative abundances of the seven CKD-associated genera were analyzed in a longitudinal cohort comprising 300 Control FF samples and paired first and second fecal samples from 150 Future CKD dogs. Differences between Control FF and the first Future CKD samples were assessed using two-sided unpaired Wilcoxon rank-sum tests. Changes between the first and second samples from the same Future CKD dogs were assessed using two-sided paired Wilcoxon signed-rank tests. Lines in the longitudinal plots connect samples obtained from the same dog. The current analyses report nominal, unadjusted P values across seven genera and two comparisons and are therefore considered exploratory. A complementary genus-wide analysis was performed after removing non-bacterial, mitochondrial and chloroplast sequences and recalculating within-sample relative abundances. Genera were eligible if they were detected in at least 5% of the 300 longitudinal samples (150 pairs) or had a combined mean relative abundance of at least 0.01%. First and second samples were compared using two-sided paired Wilcoxon signed-rank tests. Log2 fold change was calculated as log2[(mean abundance at the second sample + 1 × 10⁻⁶)/ (mean abundance at the first sample + 1 × 10⁻⁶)], with abundances expressed as percentages. P values were adjusted using the Benjamini–Hochberg method across all 99 eligible genera.

### Gut CKD dysbiosis score

The five-genus score was derived from the seven genera that met the genus-wide FDR threshold in the Future CKD–Control F comparison. Genera were retained based on prediagnostic temporal support, abundance and directional consistency. Pseudomonas was excluded because of its very low abundance and because the direction of its longitudinal change was opposite to that observed in the discovery comparison. Peptoclostridium was excluded because it did not reach the adjusted significance threshold in any individual prediagnostic interval and showed no significant change in the paired longitudinal analysis.

The resulting five-genus gut CKD score comprised Clostridium sensu stricto 1 and Incertae Sedis (positive-direction genera) and Prevotella 9, Blautia and Sutterella (negative-direction genera). For each genus, percentage relative abundance was transformed as log (1 + abundance) and standardized using the mean and standard deviation in the reference controls (Control F and Control E in the case-control implementation). The score was calculated as the mean z score of the two positive-direction genera minus the mean z score of the three negative-direction genera. Higher values therefore indicate a more CKD-like profile within this dataset. Because the component genera were selected and the score was evaluated in the same case-control cohort, the score was treated as a descriptive summary, not as a predictive model. No cross-validation or external validation was used to estimate predictive performance.

### Laboratory and microbiome trajectories before CKD diagnosis

For the 127 Future CKD dogs with linked laboratory data, BUN and creatinine were aligned to the date of the corresponding health examination, whereas microbiome measurements were aligned to the fecal sample date. Only measurements preceding the first qualifying CKD claim were included. Time before diagnosis was grouped as >730, 366-730, 181-365, 91-180 or 0-90 days. BUN and creatinine were standardized to the mean and standard deviation among prediagnostic dogs measured >730 days before diagnosis. Shannon entropy and the gut CKD score were plotted on their original scales, with matched-control means ± s.e.m. shown as reference bands. Values are presented as means ± s.e.m. Because the number of available dogs varied by analyte and time bin and the measurements were retrospectively aligned, these trajectories were interpreted descriptively.

### Paired oral-gut analysis and periodontal genus burden

ASV– and genus-level Bray-Curtis dissimilarities were calculated for the 52 paired oral and gut samples. PCoA was used to visualize site separation. Site was tested by paired PERMANOVA with dog identity used as the permutation block; permutations were restricted within dogs so that oral and gut labels were exchanged only within each matched pair. The oral periodontal burden (“Oral Perio7”) was defined as the summed oral relative abundance of Porphyromonas, Treponema, Tannerella, Fusobacterium, Prevotella, Campylobacter and Peptostreptococcus. Associations between Oral Perio7 and the gut CKD score, and between each oral periodontal genus and each score component, were assessed using two-sided Spearman rank correlations. Analyses were repeated within the predefined age strata (0-4, 5-8 and ≥9 years). Raw P values are shown for the exploratory analyses in Fig. 6 and are described as nominal because they were not adjusted for multiple comparisons; BH-adjusted q values were calculated within prespecified correlation families and are reported in Fig. 6c. Covariate-adjusted associations were evaluated using partial Spearman correlations after rank transformation and residualization for age and sex; 49 dogs with complete age and sex information were included.

### Statistics and reproducibility

Categorical characteristics in the discovery cohort were compared using chi-square tests, and continuous variables were compared using two-sided Welch t tests. Incident CKD was modelled using multivariable logistic regression including age, sex and body-size category; effects are reported as ORs with 95% CIs. For visualization of cumulative CKD incidence, we used a pre-existing classification developed by Anicom Holdings, Inc. for routine assessment of the canine gut environment before the present study. Shannon entropy was converted to a diversity deviation score using breed– and age-specific reference distributions derived from claim-free dogs belonging to the ten most frequently represented breeds in the insurance database. According to the pre-existing body-size-specific classification, dogs were categorized as having low versus high diversity using cutoffs of <43 versus ≥43 for small dogs, <45 versus ≥45 for medium-sized dogs and <48 versus ≥48 for large dogs. These thresholds had been established before the present study to identify a lower-diversity subgroup corresponding to approximately the bottom 30–45% of the score distribution and were not selected or optimized using CKD status or outcomes in the present cohort. Follow-up time was measured from the date of microbiome sampling to the first qualifying CKD claim, and the 1-year analysis was administratively limited to 365 days after sampling. Time to incident CKD was analyzed using Cox proportional-hazards models, with Shannon entropy entered as a continuous predictor in univariable, age-adjusted and age-, sex– and body-size-adjusted models. A sensitivity analysis excluded diagnoses occurring within 90 days after sample collection. The primary survival inference used continuous Shannon entropy; the low-versus-high grouping in Fig. 2b was used for visualization. Each dog was treated as an independent biological replicate except in explicitly paired longitudinal and oral-gut analyses. All tests were two-sided. Unless otherwise specified, P < 0.05 was considered statistically significant. Multiple testing was controlled using the Benjamini-Hochberg procedure within each prespecified family of comparisons. Analyses were performed in R version 4.2.3 and QIIME 2 framework version 2024.10.1, including q2-phylogeny version 2024.10.0. Exact P values, sample sizes and data underlying the figures are provided in the Source Data file.

## DATA AVAILABILITY

The data used in this study are proprietary data held by Anicom Specialty Medical Institute Inc. and are not publicly available because of commercial, confidentiality and data-governance restrictions. Requests for access to the data may be directed to the corresponding author and will be considered in consultation with Anicom Insurance, Inc., subject to applicable institutional and data-use restrictions.

## CODE AVAILABILITY

Custom scripts used for microbiome sequence processing and statistical analyses are publicly available at GitHub (repository: hitominagi/canine-ckd-microbiome-analysis, release v1.0.0). No proprietary or individual-level Anicom data are included in the repository.

## ACKNOWLEDGEMENTS

We thank the participating dog owners and veterinarians for their contributions to sample collection. We also thank Nobuaki Komori (Anicom Holdings, Inc.), Kai Ataka (Anicom Insurance, Inc.), Mizuki Inukai (Anicom Insurance, Inc.), and Hirotaka Ishida (Anicom Insurance, Inc.) for their contributions to data management, health-record curation, and support of this study. We also thank Masaya Yamaguchi (Microbial Research Center for Health and Medicine National Institutes of Biomedical Innovation, Health and Nutrition) for critically reading this manuscript. This work was partially supported by Grant-in-Aid for Challenging Exploratory Research (26K23597) and Grant-in-Aid for Scientific Research (B) (23K27789) from JSPS awarded to T.S.

## AUTHOR CONTRIBUTIONS

H.OM. and M.I. conceived the study. H.OM. performed data analysis. N.F. and K.T. performed data curation. K.T. provided and managed insurance-claims data. H.OM., M.I. and T.S. wrote the manuscript, and M.I. and T.S. supervised the study. T.S. acquired funding. All authors reviewed and approved the final manuscript.

## ADDITIONAL INFORMATION

Supplementary information will be provided as separate files.

## AI DISCLOSURE

OpenAI Codex was used to assist in developing, debugging and executing scripts for downstream data processing, statistical analyses and figure generation after ASV inference and taxonomic assignment. ChatGPT was used to assist with the evaluation and presentation of analytical results and with English-language editing of the manuscript. The authors reviewed the analytical procedures, verified the final outputs against the source data, edited all AI-assisted text and retained full responsibility for the study design, selection of the final analyses, interpretation and conclusions.

## Figure legends

**Supplementary Figure 1.** Temporal and longitudinal patterns of Pseudomonas and Peptoclostridium. **a**. Relative abundance of Pseudomonas and Peptoclostridium in Control F (n = 1,512) and Future CKD dogs stratified by time to diagnosis (>730 days, n = 159; 366-730 days, n = 273; 181-365 days, n = 167; 0-180 days, n = 157). Boxes show medians and interquartile ranges and whiskers extend to 1.5 times the interquartile range; outliers are not displayed. Comparisons with Control F used two-sided Wilcoxon rank-sum tests with BH correction across the seven genera and four-time bins. b. ASV-level decomposition of the absolute genus-level shift, as in Fig. 4c. c. Relative abundance in Control FF (n = 300) and paired first and second samples from 150 Future CKD dogs. Lines connect samples from the same dog. Tests were two-sided Wilcoxon rank-sum (Control FF versus first sample) and paired Wilcoxon signed-rank (first versus second sample). **P <0.01; n.s., not significant.

## Supplemental Tables

**Supplementary Table 1.** Most common insurance-claim reasons in deceased dogs and cats.

| Dogs |  |  | Cats |  |  |
| --- | --- | --- | --- | --- | --- |
| Rank | Diseases | (%) | Rank | Diseases | (%) |
| 1 | Neoplastic diseases | 15.3 | 1 | Cardiomyopathy | 21.6 |
| 2 | Traffic accidents | 7.5 | 2 | **Chronic kidney disease** | 15.8 |
| 3 | Vomiting, Diarrhea, Hematochezia | 7.1 | 3 | Feline infectious peritonitis | 13.5 |
| 4 | **Chronic kidney disease** | 5.9 | 4 | Lethargy and/or anorexia | 8.1 |
| 5 | Lethargy and/or anorexia | 5.0 | 5 | Neoplastic diseases | 7.2 |
| 6 | Pancreatitis | 4.5 | 6 | Pneumonia | 3.6 |
| 7 | Pneumonia | 4.2 | 7 | Valvular heart disease | 3.2 |
| 8 | Heatstroke | 3.5 | 8 | Accidental ingestion | 2.7 |
| 9 | Valvular heart disease | 3.3 | 8 | Vomiting, Diarrhea, Hematochezia | 2.7 |
| 10 | Epilepsy | 3.1 | 10 | Traffic accidents | 2.3 |

**Supplementary Table 2.** Discovery cohort overview.

|  |  | Total | non-CKD | CKD | p -value |  |
| --- | --- | --- | --- | --- | --- | --- |
| n |  | 140,025 | 138,810 | 1,215 |  |  |
| Sex | Male | 73,923 (52.8%) | 73,327 (52.8%) | 596 (49.1%) | 0.029 | * |
|  | Female | 66,085 (47.2%) | 65,466 (47.2%) | 619 (50.9%) |  |  |
|  | Unknown | 17 (0.0%) | 17 (0.0%) | 0 (0.0%) |  |  |
| Size | Small | 114,037 (81.4%) | 113,118 (81.5%) | 919 (75.6%) | 1.26x10<sup>-6</sup> | *** |
|  | Medium | 21,165 (15.1%) | 20,925 (15.1%) | 240 (19.8%) |  |  |
|  | Large | 4,823 (3.4%) | 4,767 (3.4%) | 56 (4.6%) |  |  |
| Age |  | 3.01 ± 2.75 | 2.97 ± 2.71 | 6.71 ± 4.21 | 1.71x10<sup>-155</sup> | *** |
| Shannon Entropy |  | 3.99 ± 0.72 | 3.99 ± 0.72 | 3.77 ± 0.71 | 7.70x10<sup>-26</sup> | *** |

Chronic Kidney Disease (CKD), Average ± Standard Deviation (SD)
Sex and body size were analyzed using the chi-square test. Age and Shannon entropy were analyzed using Welch's t-test.

**Supplementary Table 3.** Multivariable logistic regression for incident CKD.

| Variable | OR | 95%CI | p -value |  |
| --- | --- | --- | --- | --- |
| Age (per 1-year increase) | 1.374 | 1.339–1.410 | 2.23 × 10<sup>-130</sup> | *** |
| Body size: medium vs. small | 1.692 | 1.336–2.143 | 1.29 × 10<sup>-5</sup> | *** |
| Body size: large vs. small | 1.452 | 0.920–2.289 | 0.109 | n.s. |
| Sex: male vs. female | 0.884 | 0.728–1.074 | 0.214 | n.s. |

**Supplementary Table 4.** Pairwise PERMANOVA results for β-diversity.

**Bray-Curtis β-diversity**
| PERMANOVA | pseudo-F | p-value |  | R<sup>2</sup> |  |
| --- | --- | --- | --- | --- | --- |
| Control F vs Future CKD vs Existing CKD | 3.728 | 0.005 | ** | 0.00279 |  |
| Group | pseudo-F | p-value |  | R<sup>2</sup> | q |
| Control F vs Future CKD | 3.706 | 0.005 | ** | 0.001633 | 0.0075 ** |
| Control F vs Existing CKD | 5.244 | 0.005 | ** | 0.002738 | 0.0075 ** |
| Future CKD vs Existing CKD | 1.379 | 0.255 |  | 0.001194 | 0.255 |

**Weighted UniFrac β-diversity**
| PERMANOVA | pseudo-F | p-value |  | R<sup>2</sup> |  |
| --- | --- | --- | --- | --- | --- |
| Control F vs Future CKD vs Existing CKD | 4.282 | 0.005 | ** | 0.003203 |  |
| Group | pseudo-F | p-value |  | R<sup>2</sup> | q |
| Control F vs Future CKD | 4.997 | 0.005 | ** | 0.002201 | 0.0075 ** |
| Control F vs Existing CKD | 5.303 | 0.005 | ** | 0.002769 | 0.0075 ** |
| Future CKD vs Existing CKD | 1.342 | 0.23 |  | 0.001162 | 0.23 |

**Supplementary Table 5.** Top 20 candidate genera associated with CKD development.

| Rank | Genus | log<sub>2</sub> FC | p_value | FDR |  |
| --- | --- | --- | --- | --- | --- |
| 1 | Prevotella 9 | -0.37921 | 0.000379 | 0.034544 | * |
| 2 | Clostridium sensu stricto 1 | 0.352794 | 0.000691 | 0.034544 | * |
| 3 | Pseudomonas | 2.165512 | 0.001689 | 0.044289 | * |
| 4 | Blautia | -0.17731 | 0.002061 | 0.044289 | * |
| 5 | Peptoclostridium | -0.52646 | 0.002493 | 0.044289 | * |
| 6 | Sutterella | -0.26238 | 0.002657 | 0.044289 | * |
| 7 | Incertae Sedis | 0.404053 | 0.003184 | 0.045482 | * |
| 8 | Phascolarctobacterium | -0.41006 | 0.00624 | 0.078003 |  |
| 9 | Erysipelotrichaceae UCG-003 | -1.80789 | 0.00816 | 0.090666 |  |
| 10 | [Eubacterium] oxidoreducens group | 0.881633 | 0.010592 | 0.105922 |  |
| 11 | Clostridioides | 0.797349 | 0.013609 | 0.12372 |  |
| 12 | Parasutterella | -0.28042 | 0.015733 | 0.13111 |  |
| 13 | Escherichia-Shigella | 0.227506 | 0.021279 | 0.155634 |  |
| 14 | Flavonifractor | -0.18235 | 0.021789 | 0.155634 |  |
| 15 | [Clostridium] innocuum group | 0.128551 | 0.023616 | 0.157443 |  |
| 16 | Terrisporobacter | 0.363417 | 0.026784 | 0.167397 |  |
| 17 | Muribaculaceae | -0.12217 | 0.028729 | 0.168996 |  |
| 18 | Allobaculum | -0.24317 | 0.043035 | 0.239082 |  |
| 19 | Parabacteroides | -0.08664 | 0.050434 | 0.265443 |  |
| 20 | Slackia | -0.20924 | 0.067624 | 0.285654 |  |

**Supplementary Table 6.** Longitudinal changes in candidate gut bacterial genera.

| Rank | Taxon | log<sub>2</sub>FC | p-value | FDR |  |
| --- | --- | --- | --- | --- | --- |
| 1 | Parasutterella | −1.572 | $3.18 \times 10^{-5}$ | 0.0031 | * |
| 2 | Romboutsia | 0.777 | 0.0007 | 0.0347 | * |
| 3 | Pseudomonas | −1.915 | 0.00162 | 0.0533 |  |
| 4 | Turicibacter | 0.929 | 0.00307 | 0.0761 |  |
| 5 | Blautia | 0.411 | 0.00546 | 0.0942 |  |
| 6 | Epulopiscium | 1.883 | 0.00571 | 0.0942 |  |
| 7 | Actinomyces | 2.257 | 0.00685 | 0.0969 |  |
| 8 | Escherichia–Shigella | 0.326 | 0.011 | 0.1364 |  |
| 9 | Saccharimonadaceae | 1.85 | 0.0149 | 0.1515 |  |
| 10 | Faecalitalea | 0.383 | 0.0153 | 0.1515 |  |
| 11 | [Clostridium] innocuum group | 0.772 | 0.0181 | 0.1631 |  |
| 12 | Lachnospiraceae UCG-004 | −1.451 | 0.0224 | 0.1849 |  |

**Supplementary Table 7.** Cohort Overview of Oral-Gut cohort.

|  |  |  |
| --- | --- | --- |
| Total |  | n=52 |
| Sex | Male: Female | 25: 27 |
| Size | Small | 32 (61.5%) |
|  | Medium | 10 (19.2%) |
|  | Large | 4 (7.7%) |
|  | Unknown | 6 (11.5%) |
| Age |  | 6.76 ± 5.54 (years) |

