## Supplementary figures and images for "Gut microbiome signatures precede chronic kidney disease diagnosis in a large canine cohort"

### Supplementary Figure 1

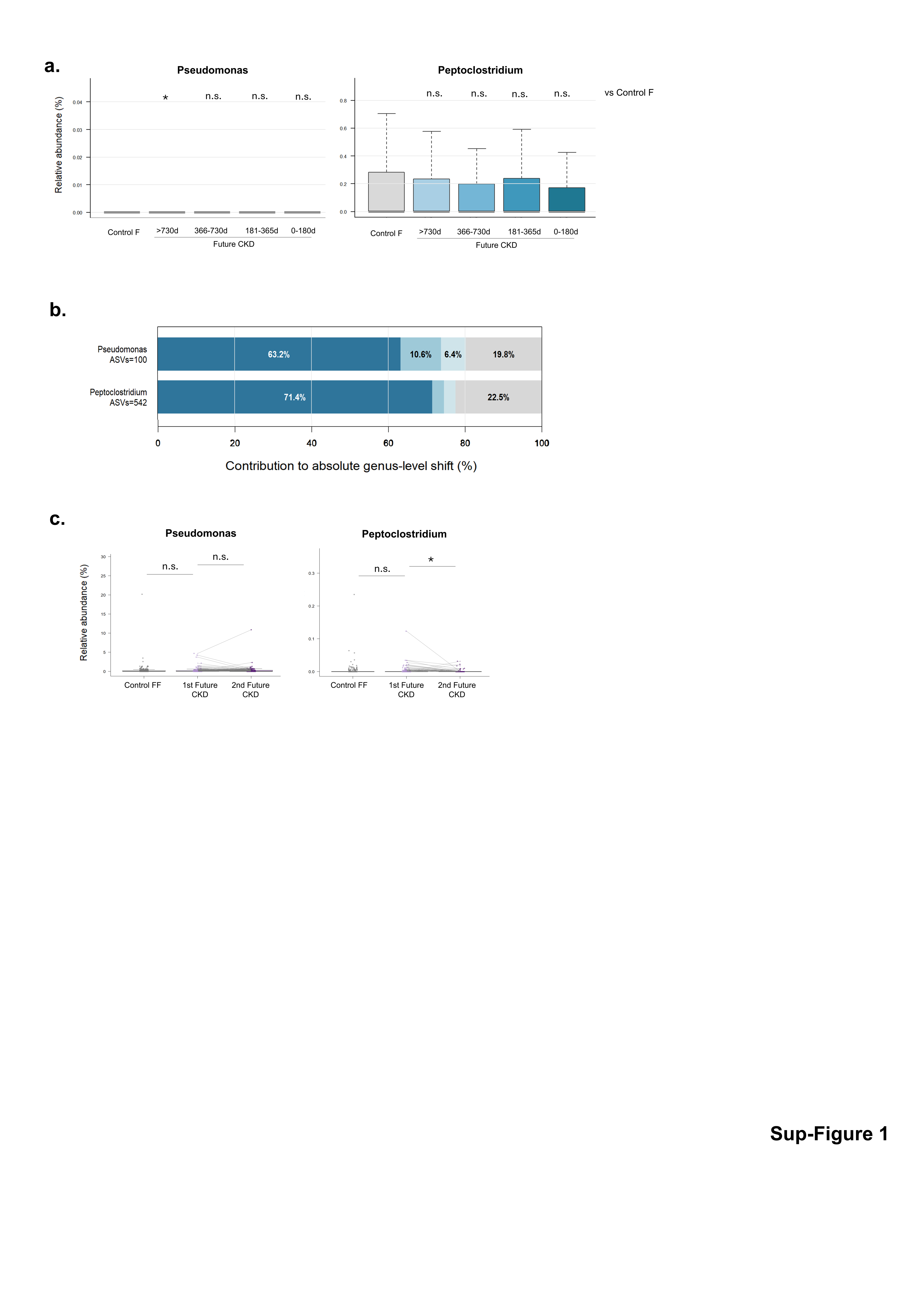
